# Metal bridge in S4 segment supports helix transition in *Shaker* channel

**DOI:** 10.1101/689620

**Authors:** Carlos A Z Bassetto, João Luis Carvalho-de-Souza, Francisco Bezanilla

**Affiliations:** Department of Biochemistry and Molecular Biology, The University of Chicago, Chicago, IL; Institute for Biophysical Dynamics, The University of Chicago, Chicago, IL; Centro Interdisciplinario de Neurociencias, Facultad de Ciencias, Universidad de Valparaiso, Valparaiso, Chile

**Keywords:** *Shaker* potassium channel, α-helix, 3_10_-helix, Metal bridge

## Abstract

Voltage-gated ion channels play important roles in physiological processes, especially in excitable cells, where they shape the action potential. In S4-based voltage sensors voltage-gated channels, a common feature is shared: the transmembrane segment 4 (S4) contains positively charged residues intercalated by hydrophobic residues. Although several advances have been made in understating how S4 moves through a hydrophobic plug upon voltage changes, possible helix transition from α-to 3_10_-helix in S4 during activation process is still unresolved. Here, we have mutated several hydrophobic residues from I360 to F370 in the S4 segment into histidine, in *i, i+3* and *i, i+6* or *i, i+4* and *i, i+7* pairs, to favor 3_10_- or α-helical conformations, respectively. We have taken advantage that His can be coordinated by Zn^+2^ to promote metal ion bridges and we have found that the histidine introduced at position 366 (L366H) can interact with the introduced histidine at position 370 (stabilizing that portion of the S4 segment in α-helical conformation). In presence of 20 μM of Zn^+2^, the activation currents of L366H:F370H channels were slowed down by a factor of 3.5, the voltage-dependence is shifted by 10 mV towards depolarized potentials with no change on the deactivation time constant. Our data supports that by stabilizing a region of the S4 segment in α-helical conformation a closed *(resting or intermediate)* state is stabilized rather than destabilizing the open (*active)* state. Taken together, our data indicates that the S4 undergoes α-helical conformation to a short-lived different secondary structure transiently before reaching the *active* state in the activation process.

**STATEMENT OF SIGNIFICANCE:** Conformational transitions between α-helix and 3_10_-helix in the S4 segment of *Shaker* potassium channel during gating has been under debate. The present study shows the coordination by Zn^2+^ of a pair of engineered histidine residues (L366H:F370H) in the intermediate region of S4 in Shaker, favoring α-helical conformation. In presence of 20μM of Zn^+2^ the activation currents of L366H:F370H channels become slower, with 10 mV positive shift in the voltage-dependence and no effects on deactivation time constants suggesting a stabilization of a closed state rather than destabilization the open *(active)* state. Collectively, our data indicate that S4 undergoes secondary structure changes, including a short-lived secondary structure transition, when S4 moves from the *resting* to the *active* state during activation.

## INTRODUCTION

Voltage-gated ion channels (VGIC) play several roles in physiological processes, especially in excitable cells, shaping the generation and propagation of action potentials (1). It has been shown by biochemical, electrophysiological and structural biology experiments that VGIC are formed by four subunits (or domains), each one containing six transmembrane (TM) segments (S1-S6) (2, 3). In each subunit, segments S1 to S4 constitute the so-called S4-based voltage sensor domain (VSD) and the segments S5 and S6 form a fourth of the pore domain. In S4-based VSDs, S4 contains positively charged residues (mainly constituted by arginine side chains), every three residues between hydrophobic residues. These charged residues sense the electric field across the membrane and move upon depolarization, conferring to the pore the ability to conduct ions (4). Segment S2 contains a negative residue (E283) that forms salt bridges with Arg (R) residues from the S4 segment stabilizing and guiding their movement (5). Also, it has been demonstrated that S1, S2 and S3 segments contain a set of hydrophobic residues that form the “hydrophobic plug”, through where S4, the side chains of the R residues, must pass to go from the *resting* to the *active* state, the latest linked to the increase in the open probability of the pore domain (6–9).

Recently, crystal structures of some VGICs have been solved and it has been noted that the S4 segments from different VGICs show different secondary structures. For example in Kv1.2, a mammalian voltage-gated potassium channel that is homologous to a potassium channel from fruit fly named *Shaker*, S4 appears in α-helical conformation in its N-terminal half while its C-terminal half exhibits 3_10_-helical conformation (7). In NavAb, a bacterial sodium channel from *Arcobacter butzleri*, the entire S4 segment is shown in 3_10_-helical conformation (10). In Ci-VSP, a VSD that connects and controls an intracellular phosphatase in *Ciona intestinalis*, the S4 segment is shown in α-helical conformation in its entirety in two structures, one near the *resting* state and the other near the *active* state (11). The Human sodium channels have four homologous domains I-IV, each domain containing six transmembrane segments similar to a potassium channel subunit. The structure of the Human sodium channel NaV1.4 has been reported, and it shows that S4 segment is organized in 3_10_-helical conformation between R2 and R6 (sixth Arg in S4) for all domains but the N-terminal region of S4 segment in domain III and IV in α-helical conformation (around R1, first Arg in S4) (12).

From a functional data perspective, by using His scanning on S4 and analyzing periodicity of the perturbation using gating currents (13), and by using metal-ion bridges for putative intersegment interactions between S3 and S4 (8), it has been suggested that S4 from *Shaker* can adopt a transient 3_10_-helical conformation during the activation process. Nonetheless, a direct physical measurement of the interconversion from α to 3_10_-helical secondary structure has not yet been determined for S4. An attempt to probe 3_10_ conversion from alpha structure in Ci-VSP showed that the conversion is not required in this voltage sensor (14). Recently, using nonsense suppression to investigate the hydrogen bonding of the main-chain of *Shaker* S4 segment, Infield et al (2018) have found that the *active* conformation of segment S4 was destabilized when the hydrogen bond in residue 369 was perturbed. This data suggests that destabilization of the helical conformation interferes with the movement of S4 segment from *resting* to the *active* state. Even though those reports have suggested conformational transition in different regions when S4 segment reaches the *active* state (8, 13), no other direct evidence of a conformational transition has been provided, which keeps the matter under debate.

Metal ion bridges and disulfide bonds have been widely used to probe movement and distances between different residues in VGIC proteins (8, 15–18, 18–21). This method has been especially used to probe interactions between residues in segment S4 and S5 in *Shaker* (16) and K_V_1.2 (18), as well as between segments S3 and S4 in *Shaker* (8, 19). We engineered a pair of histidine residues within the S4 segment for coordination with Zn^2+^ to determine transitions from α-to 3_10_-helical conformation or vice-versa in this transmembrane segment. We placed His residues pairs in different positions to favor one or the other helical conformation. In two cases (*i,i+3* and *i,i+6*, where *i* is a residue number), the His-His pair are expected to be aligned in 3_10_-helical structures and would stabilize the helix in this conformation when coordinated by Zn^2+^. In other two cases (*i,i+4* and *i,i+7*), Zn^2+^ is expected to coordinate the His-His pair in α-helical structures, stabilizing it. We have found strong evidence that the histidine introduced at position 366 (L366H) coordinates with the introduced histidine at position 370, therefore favoring the α-helical conformation (*i,i+4*) in presence of μM Zn^+2^ concentrations The kinetics of the activation of K^+^ currents in L366H:F370H mutant channels were slowed down by a factor of 3.5, the voltage-dependence was rightward shifted (about 10 mV) and deactivation time constant was unchanged when 20 μM of Zn^+2^ was present in external solution. These results imply that the stabilization of a region of the S4 segment in α-helical conformation counteracts the transition of VSD to its *active* state. Taken together, our data supports the evidence that the S4 undergoes a secondary structural transition from α-helical to a short-lived secondary structural conformation, different from α-helical, when transiting from the *resting* state to the *active* state during the activation process.

## MATERIALS AND METHODS

### Site-directed mutagenesis

We used *Shaker* zH4 K^+^ channel with fast inactivation removed by deleting residues from 6 to 46, Δ6-46, cloned into pBSTA vector (22). Mutations were performed using Quick-change (Stratagene, La Jolla, CA). All plasmids containing wild type or mutant *Shaker* cDNA were sequenced and then linearized by restriction enzyme NotI. cRNA was transcribed using *in vitro* transcription kits (T7 RNA expression kit; Ambion Invitrogen). After 12-24 hours from harvesting and defoliculation, oocytes stage V-VI were injected with cRNA (5 to 100 ng diluted in 50 nl of RNAse free water). Injected oocytes were incubated from 1 to 3 days at 12°C or 18°C, depending on the construct, in standard oocytes solution (SOS) that contains the following components in mM: 100 NaCl, 5 KCl, 2 CaCl_2_, 0.1 Ethylenediaminetetraacetic acid (EDTA) and 10 4-(2-Hydroxyethyl)piperazine-1-ethanesulfonic acid, N-(2-Hydroxyethyl)piperazine-N′-(2-ethanesulfonic acid) HEPES – the pH was set to 7.4. SOS was supplemented with 50 μg/ml gentamycin to avoid contamination during incubation.

The 3_10_- and α-helix are distinct in their backbone hydrogen bonding pattern. In 3_10_-helices, the carbonyl group of a residue *i* binds to the nitrogen from amide group in residue *i+3*. In α-helices, the residue *i* from carbonyl group binds to the nitrogen from amide group in residue *i+4*. Common features of canonical 3_10_-helices are: 3 residues per turn, the angle between the residues is 120° around the helical axis, and each turn is spaced by 6 Å. For α-helices the common features are: 3.6 residues per turn, the angle between the residues is 100° around the helical axis, and each turn is spaced by 5.4 Å. Thus side chains of coupled residues spaced in *i, i+4* and *i, i+7* patterns tend to stabilize the α-helical conformation because they are closer to multiples of 3.6, and the angle between *i* and *i+4* is 400° and between *i* and *i+7* is 700°. On the other hand, side chains spaced in *i, i+3* and *i, i+6* patterns tend to stabilize the 3_10_-helical conformation since the angle between residues *i* and *i+3* is 360° and between *i* and *i+6* is 720°. Considering His:His mutants arranged in *i, i+4* and *i, i+7*, the side chains would be in opposite directions if S4 were in a 3_10_-helical conformation, being an impediment for a side chain interaction. In contrast, His:His mutants arranged in *i, i+3* and *i, i+6*, the side chains would also be in opposite directions if S4 were in an α-helical conformation. Therefore the strategy of His:His mutants was intended to favor one or the other helical conformation depending on the position where they are placed.

### Electrophysiological recordings

Ionic currents were recorded using cut-open oocyte voltage-clamp (COVC) method with a voltage-measuring pipette with resistance from 0.2 to 0.8 MΩ (23). Pipettes were pulled using a horizontal puller (P-87 Model-Sutter Instruments, Novato, CA). Capacitive transient currents were compensated by a dedicated circuit. Raw currents were analogically filtered at 20-50 kHz with a low pass 4-pole Bessel filter built-in the amplifier (Dagan CA-1B, Dagan, Minneapolis, MN). Processed currents were then sampled at 1 MHz using a 16-bit analog-to-digital converter (USB-1604, Measurement Computing, Norton, MA), digitally filtered with Nyquist frequency and decimated for an acquisition rate of 100-200 kHz. An in-house software was used to acquire (GPatch64MC) and analyze the data (Analysis). For ionic measurements the external solution was composed by (mM): K-MES – MES (Methanesulfonic acid), 12, Ca-MES 2, HEPES 10, EDTA 0.1, NMDG-MES – NMDG (N-Methyl-D-glucamine) 108, pH 7.4 and the internal solution was composed by (mM): K-MES 120, EGTA 2, HEPES 10, pH7.4. To ensure the channels were closed before the depolarizing pulses, a conditioning hyperpolarized pre-pulse was used with different voltages (depending on the mutant) and the holding potential used was −80 mV or −100 mV depending on the mutants. All recordings were performed at room temperature. Prior to use, Zn^2+^ was diluted to its final concentration in external solution in absence of EDTA, from a 30 mM stock solution (ZnCl_2_).

### Data analysis

Macroscopic K^+^ currents were recorded from four to eleven oocytes per mutant and their peak was transformed into conductance (*G*) by using the equation below:

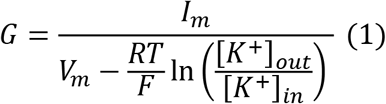

where *I_m_* is the K^+^ current activated by membrane voltage *V_m_, R* is the gas constant, *T* is the temperature in Kelvin, *F* is the Faraday constant, and [*K*]_*in*_ and [*K*]_*out*_ are the intracellular and extracellular K^+^ concentrations, respectively.

The K^+^ conductances were normalized, averaged and plotted against *V_m_* in order to build conductance-voltage (G-V) curves that were fitted with a two-state model by the equation below:

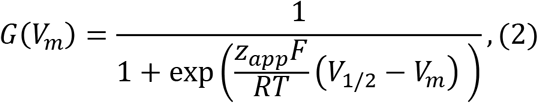

where *z_app_* is the apparent charge of the transition expressed in units of elementary charge (*e*_0_) and *V*_1/2_ is the voltage for 50% of the maximal conductance

K^+^ currents during activation by depolarizing voltage pulses and during deactivation by hyperpolarizing voltage pulses were fitted by a double exponential shown below:

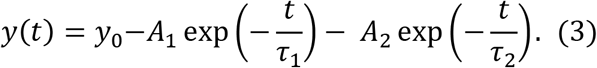

where *A*_1_ and *A*_2_ are the amplitudes from the first and the second exponential, respectively. *τ*_1_ and *τ*_2_ are the time constants for the first and second exponentials, respectively; *y*_0_ is a baseline adjustment.

For analysis purposes, we considered a weighted time constant, calculated by the following equation:

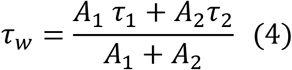

where *τ_w_* is the weighted time constant.

Matlab (The MathWorks, Inc., Natick, MA) and Origin9.0 (Origin Lab Corporation, Northampton, MA) were used for calculating G-Vs, τ-Vs, plotting and fitting the data. Data is shown as Mean ± standard error of mean (SEM).

## RESULTS

We tested the hypothesis whether S4 segment of *Shaker* undergoes secondary conformational transitions during the activation/deactivation processes. To that end, we replaced pairs of non-charged residues located at different positions of S4 helix by pairs of His (Fig. 1 A-B). Each pair of residues replaced by His were chosen so that the His:His coordination by Zn^2+^ would be favored exclusively in α-helical or 3_10_-helical conformations of the secondary structure of S4 region (Fig. 1 A). For example, when a His:His pair was placed every four or seven residues apart (*i, i+4* or *i, i+7*), a possible interaction between them would favor an α-helical conformation. On the other hand, when they were placed every three or six residues apart (*i, i+3* or *i, i+*6) they would favor a 3_10_-helical conformation (See methods). His:His were introduced in the fast inactivation removed version of the *Shaker* channel (Δ6-46, *Shaker* IR) (22). We analyzed possible His:His interactions using the voltage-dependence of the K^+^ conductance (G-V curves) as well as the kinetics of activation and deactivation of the voltage-dependent K^+^ currents. If a conformation change were to occur during activation or deactivation, the possible stabilization of either secondary structure conformation by a His:His interaction may oppose or promote that process, possibly altering the activation/deactivation kinetics and/or voltage-dependence. To enhance or disrupt His:His interaction we exposed the mutant channels to different pHs. Acidic environment at pH 5.5 would protonate the majority of the exposed His residues whereas basic environment at pH 9.2 would deprotonate the majority of the exposed His residues. Zn^+2^ was also used to coordinate the His:His by metal bridges (16, 24, 25).

**Figure 1:**
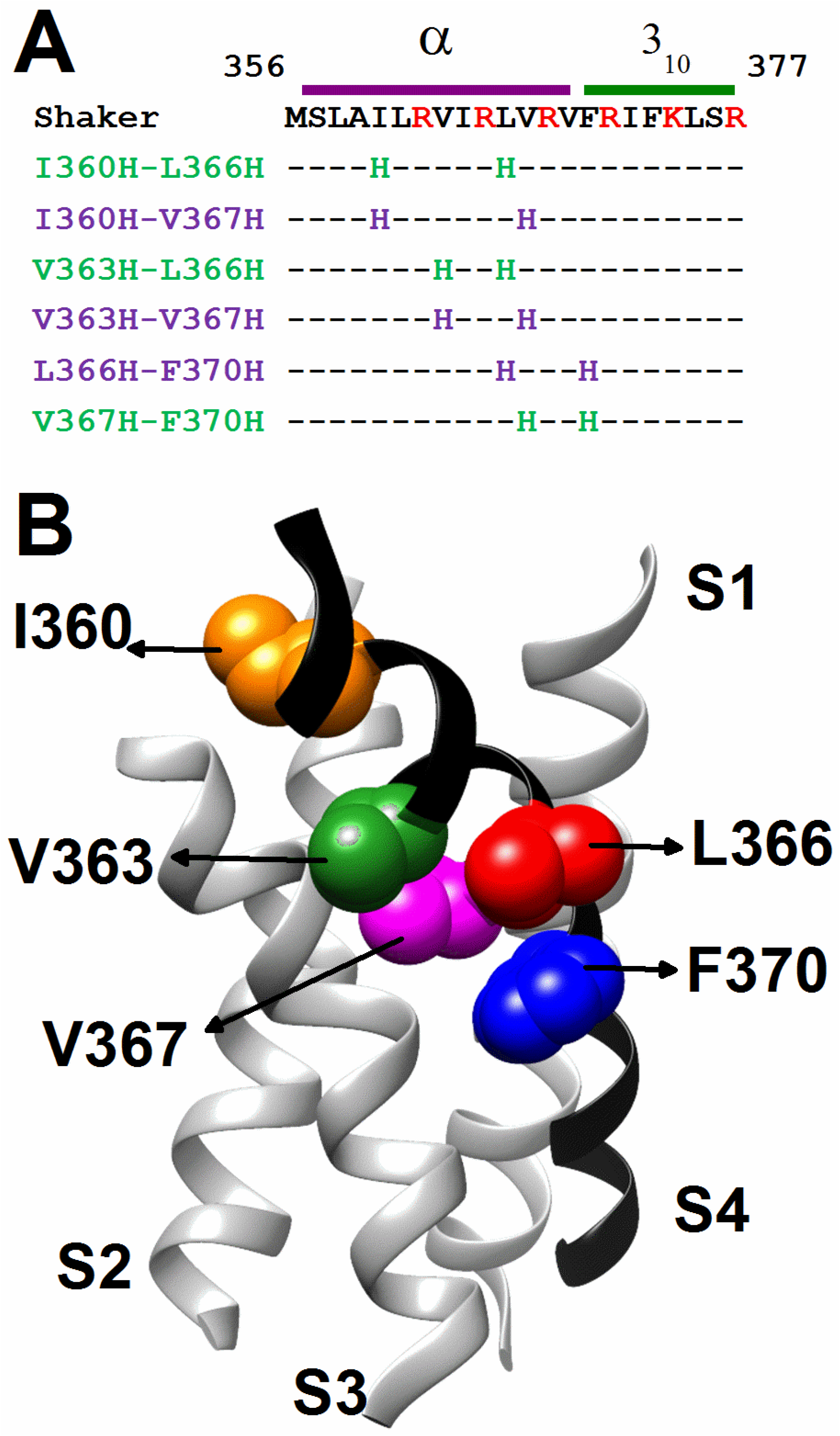
His:His screening in S4 segment. **(A)** His:His residues mutated in S4. The top row in black shows the wild type amino acid *Shaker* S4 sequence. Arg residues are shown in red. Colored rows show the residues mutated and the dashes mean wild type residues. Rows in purple represent the mutants intended to stabilize S4 in α-helical conformation, whereas rows in green represent the mutants intended to stabilize S4 in 3_10_-helical conformation. Green and purple top bars represent the portion of S4 segment that are organized in α-helical or 3_10_-helical configuration according to K_V_1.2 VSD from a crystallography-based model (PDB code 3LUT), respectively. **(B)** Side view from S1 to S4 segments at the *active/relaxed* state in the K_V_1.2 VSD from a crystallography-based model (PDB code 3LUT). For clarity purposes, helices backbones from S1 to S3 are shown in grey and S4 in black. Residues mutated I360H (orange), V363H (green), L366H (red), V367H (magenta) and F370 (blue) are depicted with spheres.

### Zn^+2^ affects K^+^ currents of L366H:F370H channel: possible stabilization of α-helical conformation

Remarkably, the only double mutant that showed evidence of His:His interactions, as measured by changes in voltage-dependence and kinetics, was L366H:F370H in presence of 20 μM Zn^+2^. Fig. 2 A and B show representative ionic current traces from L366H:F370H mutant in presence (Fig. 2B) and absence of 20 μM Zn^+2^ (Fig. 2A). The Zn^+2^ effect on L366H:F370H was partially recovered after washout with external solution (containing 100 μM of EDTA to chelate free Zn^2+^) (Fig. 2C). In presence of Zn^+2^, the midpoint of L366H:F370H G-V curve is shifted ~10 mV to more positive potentials (from −11.4 ± 1.03 to −0.3 ± 1.5 mV) without changing the slope of the curve (from 1.4 ± 0.1 to 1.2 ± 0.1) and the maximum conductance is decreased to ~60% of the conductance of No Zn^+2^ condition (Fig. 2D – Table 1). After extensive washing, the maximum conductance was recovered to 84% of the initial value (No Zn^+2^ condition). This is clearly different from the results of Zn^+2^ on WT, or the single mutants L366H and F370H, where no effects were observed (Fig. 2E-G). These findings suggest that the coordination between L366H and F370H by Zn^2+^ is enough to oppose the activation process, indicating that a change in secondary structure conformation might be needed for that process.

**Figure 2.**
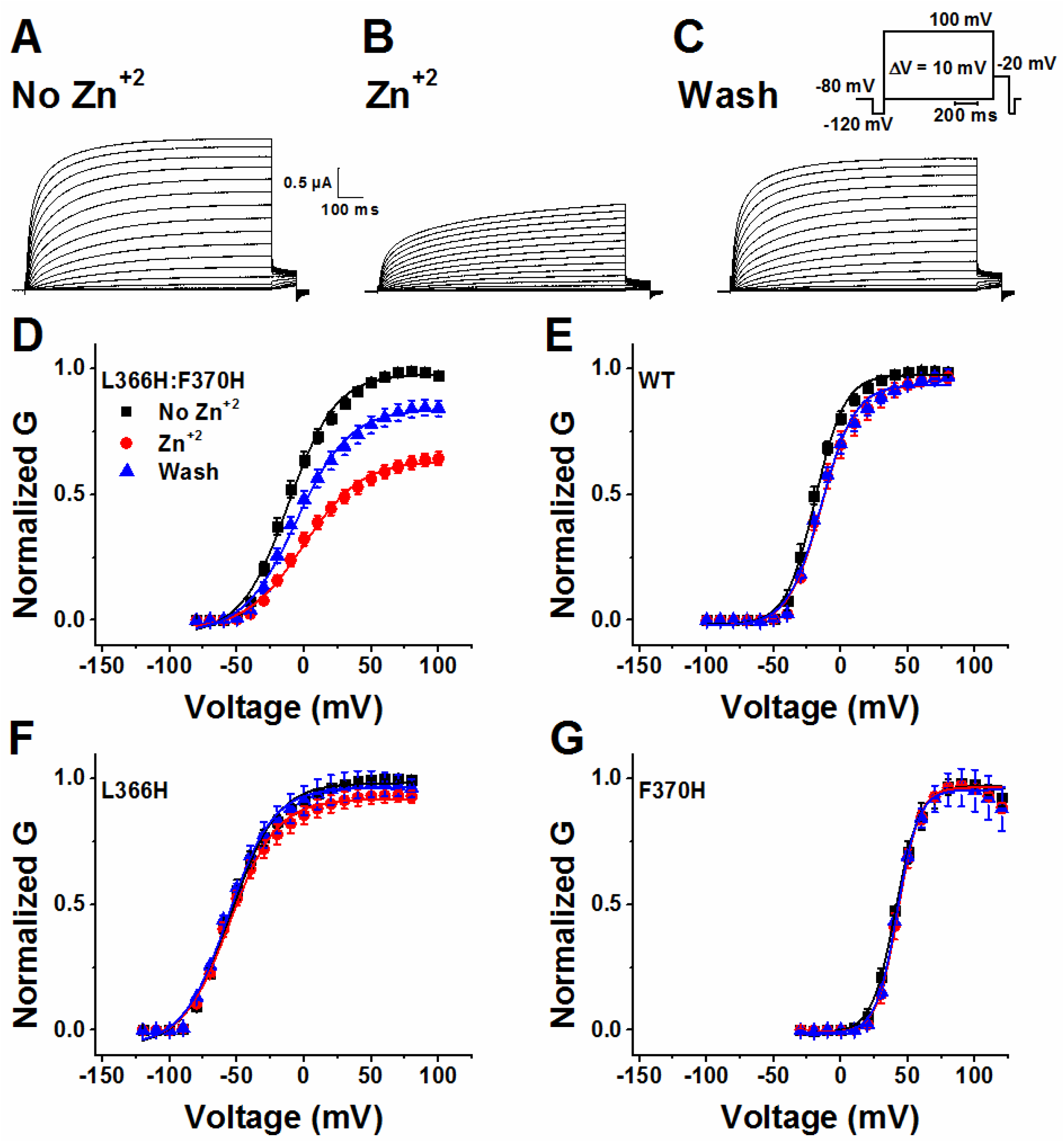
His:His coordination by Zn^+2^ between L366H and F370H in S4 segment. **(A-C)** Representative ionic currents for mutant L366H:F370H in external solution only (**A**), in presence of 20μM Zn^+2^ **(B)** and washout (external solution) **(C)**. Inset in C is the voltage pulse protocol used to elicit the activation currents. **(D-G)** G-V curves for L366H:F370H **(D)**, for WT **(E)**, for L366H **(F)** and for F370H **(G),** in absence of Zn^+2^ (No Zn^+2^, external solution – black squares), in presence of 20μM Zn^+2^ (Zn^+2^, red squares) and washout condition (Wash, external solution – blue triangles). G-Vs curves were normalized by the maximum G in absence of Zn^+2^. Data are shown as Mean ± SEM (*N* = 4-11). Continuous lines over G-V curves are the best fittings of Eq. 2 (see Table 1 for fitted parameters).

**Table 1:** G-V fitted values for all the channels at different conditions.

| Channel | pH 7.4 (No Zn<sup>+2</sup>) | | | | | | pH 5.5 | | | | | | pH 9.2 | | | | | | 20 μM Zn<sup>+2</sup> | | | | | |
| --- | --- | --- | --- | --- | --- | --- | --- | --- | --- | --- | --- | --- | --- | --- | --- | --- | --- | --- | --- | --- | --- | --- | --- | --- |
|  | V<sub>1/2</sub> (mV) |  |  | z |  |  | V<sub>1/2</sub> (mV) |  |  | z |  |  | V<sub>1/2</sub> (mV) |  |  | z |  |  | V<sub>1/2</sub> (mV) |  |  | z |  |  |
| WT | -19.1 | ± | 0.7 | 2.3 | ± | 0.1 | 4.8 | ± | 0.9 | 2.3 | ± | 0.2 | -22.9 | ± | 1.1 | 2.1 | ± | 0.2 | -14.2 | ± | 1.1 | 2.1 | ± | 0.2 |
| I360H | -4.4 | ± | 0.5 | 2.0 | ± | 0.1 | 19.3 | ± | 0.8 | 1.8 | ± | 0.1 | -8.0 | ± | 0.7 | 1.8 | ± | 0.1 | -1.9 | ± | 0.5 | 2.0 | ± | 0.1 |
| V363H | -0.4 | ± | 0.9 | 1.0 | ± | 0.0 | 29.8 | ± | 1.2 | 1.0 | ± | 0.0 | -7.6 | ± | 1.3 | 0.9 | ± | 0.0 | -0.4 | ± | 0.9 | 1.0 | ± | 0.0 |
| L366H | -54.4 | ± | 1.4 | 1.5 | ± | 0.1 | -24.3 | ± | 0.6 | 1.6 | ± | 0.1 | -63.8 | ± | 1.1 | 1.5 | ± | 0.1 | -55.2 | ± | 1.4 | 1.4 | ± | 0.1 |
| V367H | 0.0 | ± | 0.6 | 3.0 | ± | 0.2 | 30.4 | ± | 0.6 | 2.8 | ± | 0.2 | -8.7 | ± | 0.6 | 2.9 | ± | 0.2 | 2.2 | ± | 0.6 | 2.8 | ± | 0.2 |
| F370H | 40.5 | ± | 0.4 | 3.4 | ± | 0.2 | 62.2 | ± | 1.0 | 3.2 | ± | 0.2 | 35.7 | ± | 0.6 | 3.7 | ± | 0.2 | 44.8 | ± | 1.1 | 4.1 | ± | 0.3 |
| I360H:L366H | -32.1 | ± | 1.2 | 1.9 | ± | 0.1 | -12.8 | ± | 1.1 | 1.6 | ± | 0.1 | -39.5 | ± | 1.2 | 2.0 | ± | 0.2 | -26.8 | ± | 1.5 | 1.7 | ± | 0.1 |
| I360H:V367H | -2.3 | ± | 0.7 | 3.6 | ± | 0.3 | 28.7 | ± | 0.5 | 3.4 | ± | 0.2 | -12.6 | ± | 1.2 | 3.1 | ± | 0.4 | -1.4 | ± | 0.7 | 3.4 | ± | 0.3 |
| V363H:L366H | -48.2 | ± | 1.0 | 1.1 | ± | 0.0 | -19.7 | ± | 1.1 | 0.9 | ± | 0.0 | -57.8 | ± | 1.1 | 1.1 | ± | 0.0 | -42.0 | ± | 1.2 | 0.9 | ± | 0.0 |
| V363H:V367H | -4.2 | ± | 0.8 | 1.7 | ± | 0.1 | 18.6 | ± | 0.7 | 1.5 | ± | 0.1 | -12.7 | ± | 1.0 | 1.7 | ± | 0.1 | -1.3 | ± | 1.0 | 1.6 | ± | 0.1 |
| L366H:F370H | -11.7 | ± | 1.3 | 1.4 | ± | 0.1 | 18.3 | ± | 0.8 | 1.5 | ± | 0.1 | -13.7 | ± | 1.5 | 1.3 | ± | 0.1 | -0.3 | ± | 1.5 | 1.2 | ± | 0.1 |
| V367H:F370H | 38.9 | ± | 0.5 | 4.6 | ± | 0.4 | 56.2 | ± | 0.2 | 4.2 | ± | 0.1 | 35.3 | ± | 0.5 | 4.7 | ± | 0.4 | 40.5 | ± | 0.4 | 4.1 | ± | 0.2 |

All other double mutants were not clearly affected by either the presence of Zn^+2^ or by changes in the pH compared to the effect of these changes in WT channels (Fig. S1 and S2). The observed effects induced either by the presence of Zn^2+^ or pH changes were seen as modifications in the voltage dependence of G-V curves or in the activation time constants of K^+^ currents (Fig. S1 and S2 in the Supporting Material). Even though the maximum conductance of L366H:F370H channels is affected by pH5.5 (decreased to 60% Fig. S1K), we could not infer whether the His:His were interacting in L366H:F370H channels at pH 5.5 because i) the rightward shift on the G-Vs induced by pH 5.5 on WT, L366H, F370H and L366H:F370H channels were very similar (~25-30 mV Fig. S1 A, D, F K and Table 1); ii) the activation time constants for WT, L366H, L366H:F370H were similarly slowed down by the same factor: 1.3 but in the single mutant F370H the time constants were slowed down by a factor of 4 (Fig.. S2 A, D, F and K). These results indicate that the effects of pH 5.5 on L366H:F370H could not be clearly distinguished from the effects on L366H, F370H and WT and for this reason we did not further pursue the pH effects.

### Zn^+2^ metal bridge in L366H:F370H suggests stabilization of closed states

To gain more insight about L366H:F370H interaction in presence of Zn^+2^, the activation time constants were estimated by fitting a double exponential function (eq. 3) to the rising phase of the ionic currents elicited by depolarizing membrane voltage steps (Fig. 2A-C). Next, those time constants were weighted (eq. 4) and plotted against membrane voltage (Fig. 3). Interestingly, in presence of 20 μM of Zn^+2^, L366H:F370H activation currents become ~3.5 slower than the currents in absence of Zn^+2^ (Fig. 3A). The activation time constants of WT, and on the single mutants L366H and F370H channels in presence of 20 μM of Zn^+2^ are on average only ~1.3 slower than time constants in absence of Zn^+2^ (Fig. 3B-D). Taken together, the hypothetical stabilization by Zn^+2^ of an α-helical conformation between residues L366H and F370H may suggest that the transition from *resting* to *active* states in S4 segment of the channel has been delayed, presumably stabilizing the closed (*resting or intermediate)* state or destabilizing the *active* state of the channel.

**Figure 3.**
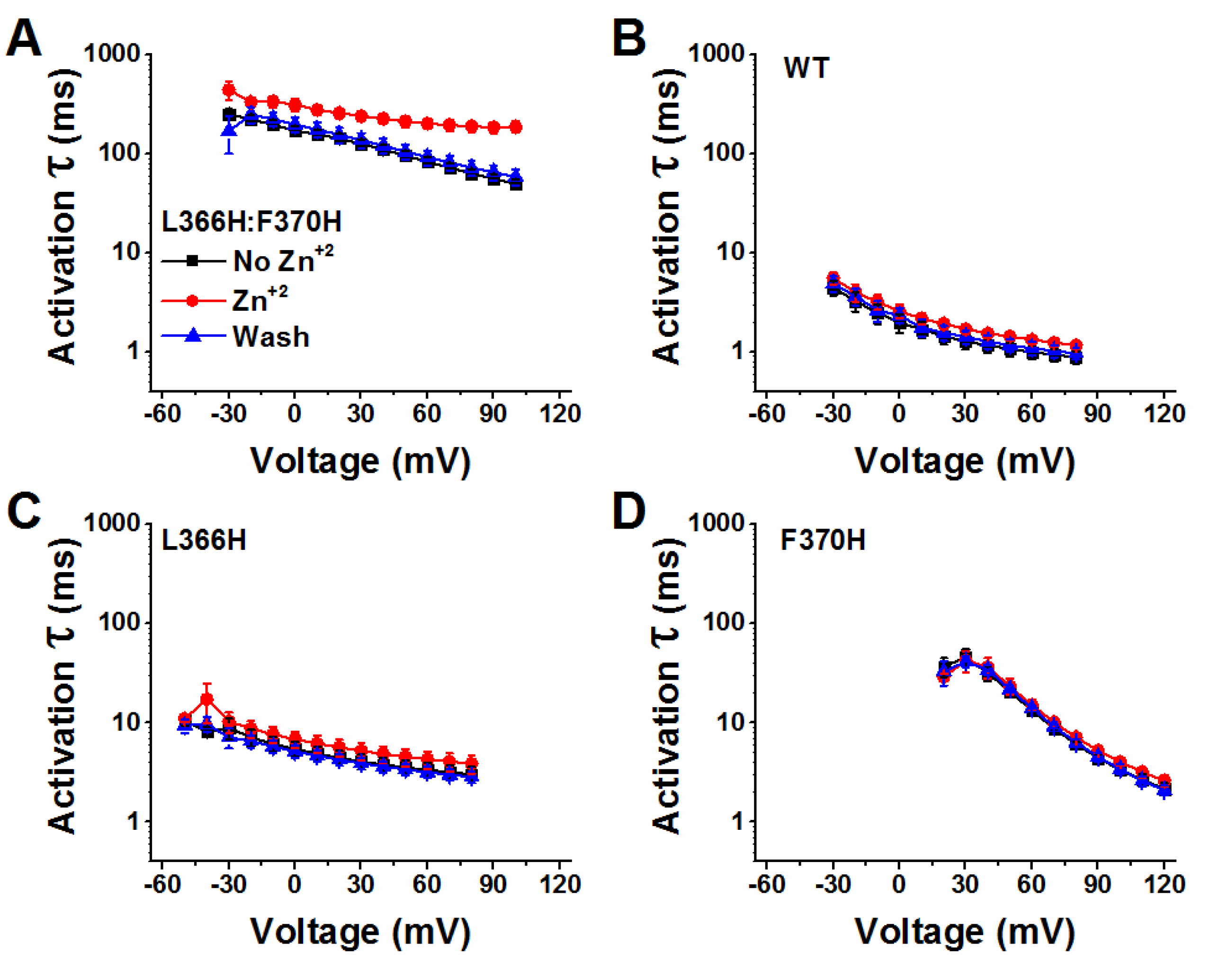
Zn^+2^ slows down activation time constants for L366H:F370H channel.) Activation τ-V curves in absence of Zn^+2^ (No Zn^+2^, external solution – black squares), in presence of 20μM Zn^+2^ (Zn^+2^, red squares) and washout condition (Wash, external solution – blue triangles). **(A)**, τ-V curves for L366H:F370H **(B)**, τ-V curves for WT **(C)** τ-V curves for L366H **(D),** τ-V curves for F370H Time constants were calculated using eq. 3 and posteriorly weighted using eq. 4. Data are shown as Mean ± SEM (*N* = 4-11).

Because the activation time constants were fitted by the sum of two exponentials (eq. 3), the analysis resulted in a fast and a slow components. We further investigate the effects of Zn^+2^ in each component. Fig. 4A shows two normalized representative ionic current traces for L366H:F370H mutant at 100mV in absence (black trace) and in presence of Zn^+2^ 20 μM (red trace). Zn^+2^ affects mainly the slow component, making it even slower (Fig. 4B) and slightly increasing its proportion with respect to the fast component (Fig. 4C). In addition, the slow component becomes fairly voltage independent in the presence of Zn^2+^, suggesting a limiting step introduced by the putative His:His coordination.

**Figure 4.**
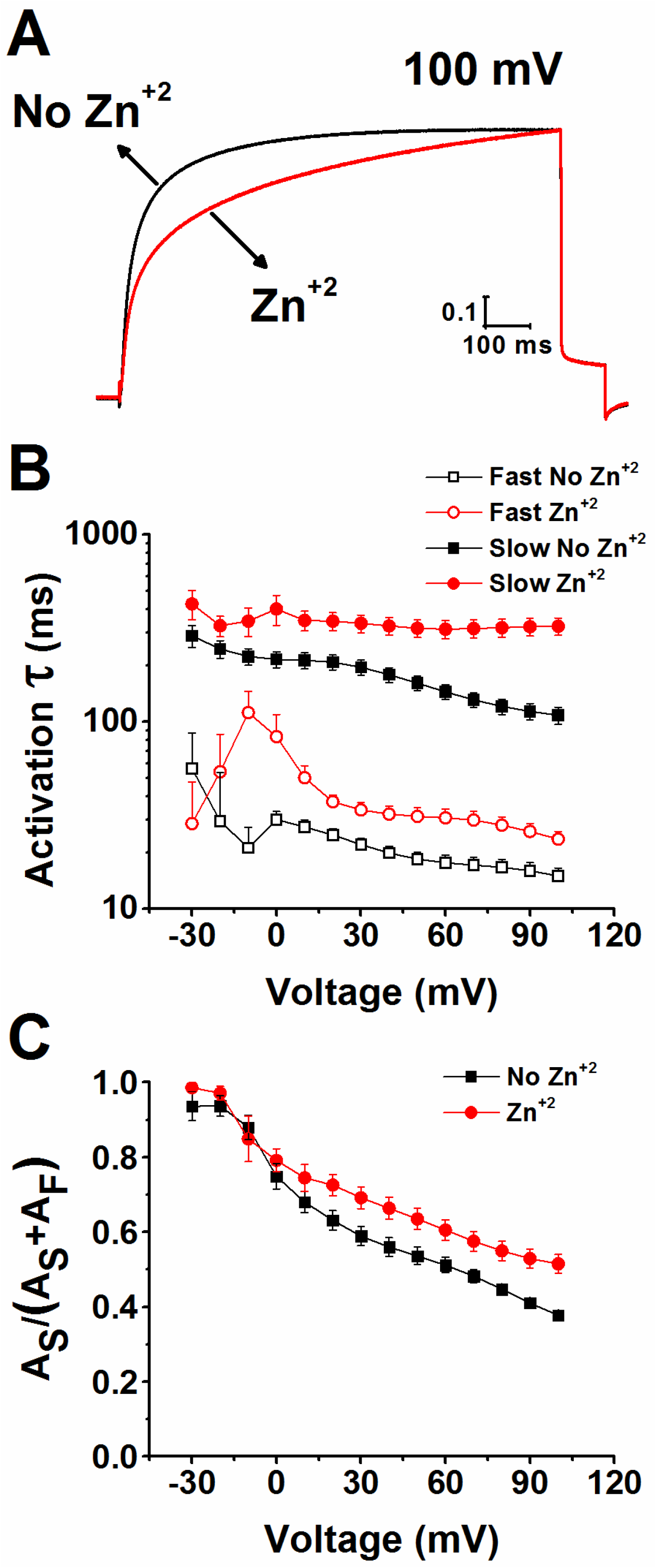
Slow component of L366H:F370H ionic currents is more affected by Zn^+2^. **(A)** Normalized representative ionic currents from an oocyte for mutant L366H:F370H at 100mV in external solution (No Zn^+2^ - black line), and in presence of 20μM Zn^+2^ (red line). **(B)** Fast (open black square – No Zn^+2^, and open red circle - 20μM Zn^+2^-Zn^+2^ condition) and slow (filled black square – No Zn^+2^ and filled red circle - 20μM Zn^+2^ - Zn^+2^ condition) component of activation time constant curves. **C)** Fraction of the slow component in the activation time constant in absence (filled black square) and in presence of 20μM Zn^+2^ (filled red square).

After 1s at positive membrane voltage the currents of L366H:F370H in presence of Zn^+2^ have not reached a steady-state and are decreased as compared to L366H:F370H currents in absence of Zn^+2^ (Fig. 2A-D). Therefore, we decided to extend the depolarizing voltage pulse to 5 s and elicit L366H:F370H currents at different Zn^+2^ concentrations in order to titrate its effect. Fig. 5A, 5B, 5C and 5D show representative L366H:F370 current traces in absence and in presence of 5, 20 and 50 μM Zn^+2^, respectively. The higher the Zn^+2^ concentration in the bath the larger is the effect on the currents (the current time courses become slower and the amplitudes smaller). The G-V is more rightward shifted (Fig. 5E-F), the proportion of the second component as well as the activation time constant are increased as the concentration of Zn^+2^ is increased in the bath (Fig. 5G-H). These results indicate that the range of Zn^2+^ concentrations we have used (5-50 μM) is not enough to saturate its binding sites in the channels (His:His), an important information for modelling this effect.

**Figure 5.**
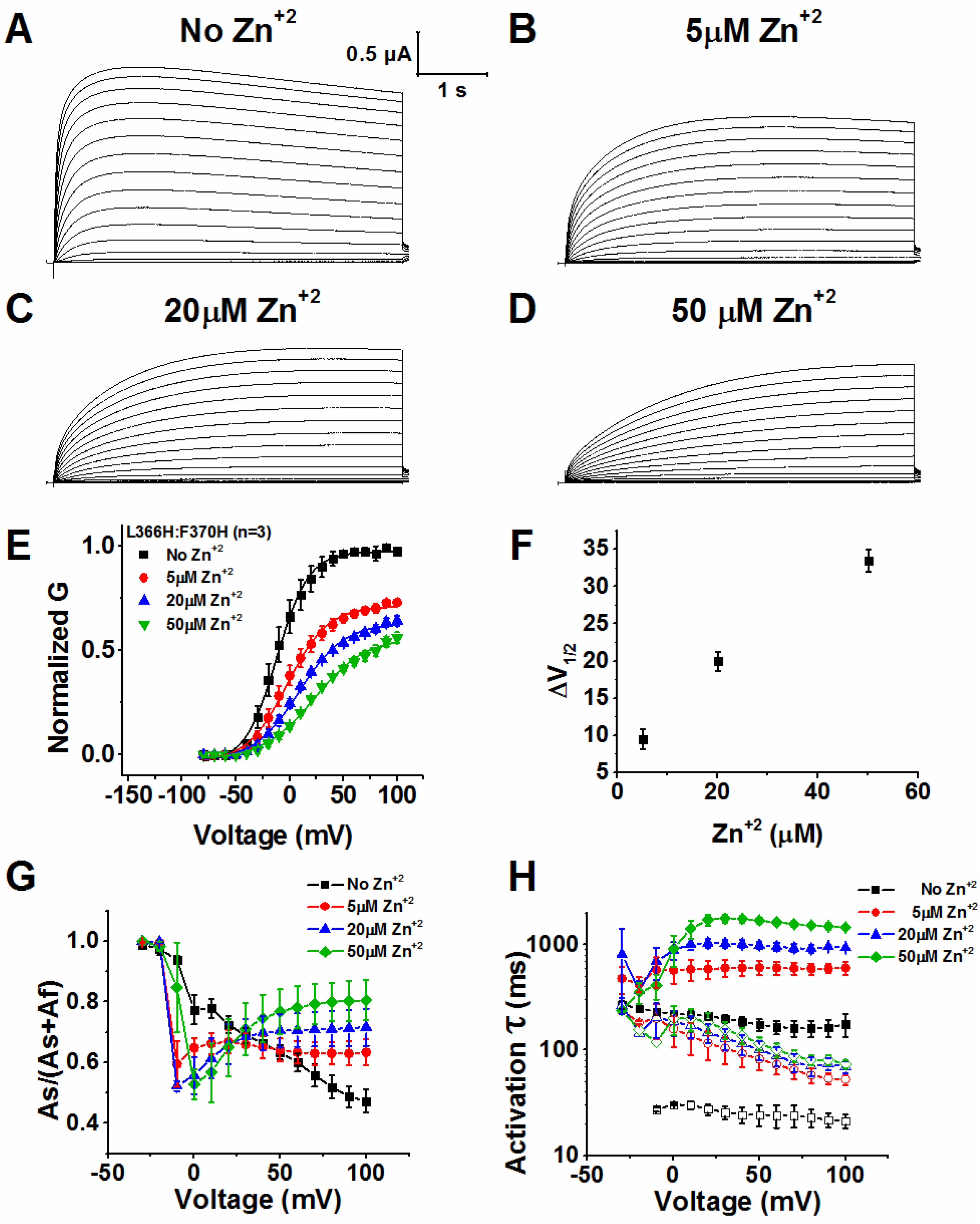
Concentration dependent effects of Zn^+2^ on L366H:F370H ionic currents. **(A-D)** Representative ionic currents for mutant L366H:F370H in external solution (**A**), in presence of 5 μM Zn^+2^ **(B)** in presence of 20 μM Zn^+2^ **(C)** and in presence of 50 μM Zn^+2^ **(D). (E)** G-V curves for L366H:F370H in absence of Zn^+2^ (No Zn^+2^ – black squares), in presence of 5 μM Zn^+2^ (red circles), 20 μM Zn^+2^ (blue triangles) and 50 μM Zn^+2^ (green triangles). G-V curves were normalized by the maximum conductance (G) of the channel in absence of Zn^+2^. Continuous lines over G-V curves are the best fittings of Eq. 2. The fitted parameters are V_1/2_= −11.9 ± 0.9 mV and z= 1.7 ± 0.1 in absence of Zn^+2^; V_1/2_= −2.2 ± 1.3 mV and z= 1.34 ± 0.1 for 5 μM Zn^+2^; V_1/2_= 8.7 ± 1.2 mV and z= 1.21 ± 0.1 for 20 μM Zn^+2^; V_1/2_= 22.8 ± 1.6 mV and z= 1.1 ± 0.1 for 50 μM Zn^+2^. **F)** ΔV_1/2_ plotted against concentration. ΔV_1/2_ was calculated using the equation: *ΔV*_1/2_ = *ΔV*_1/2_*Zn*_ – *ΔV*_1/2_*No Zn*_. **G)** Fraction of the slow component in the activation time constant in F370H in absence of Zn^+2^ (No Zn^+2^ – black squares), in presence of 5 μM Zn^+2^ (red circles), 20 μM Zn^+2^ (blue triangles) and 50 μM Zn^+2^ (green diamonds). **(H)** Fast (open symbols) and slow (filled symbols component of activation time constant for L366H:F370H in absence of Zn^+2^ (No Zn^+2^ – black squares), in presence of 5 μM Zn^+2^ (red circles), 20 μM Zn^+2^ (blue triangles) and 50 μM Zn^+2^ (green diamonds). Data are shown as Mean ± SEM (*N* = 3).

Metal bridges can stabilize the closed state or the open state of the channel depending on the position of the residues in the channel (16). Because Zn^+2^ promotes rightward shift in G-V curve and slows down the activation time constant for L366H:F370H, two possible mechanism can be speculated: i) in presence of Zn^+2^ the *resting* state is stabilized, or ii) the *active* state is destabilized. To discriminate between these possibilities, we measured the deactivation time constants of ionic currents for L366H:F370H mutant in presence (Fig. 6B) and absence of Zn^+2^ (Fig. 6A). Inset in B is the voltage pulse protocol used to record K^+^ currents during deactivation by voltage. Two normalized representative ionic currents deactivating at different voltages −70 mV (left panel) and −150 mV (right panel) in presence (red) and absence (black traces) of Zn^+2^ are shown in Fig. 6C. Interestingly, the time course of those currents practically superimposes, showing that deactivation is not affected by Zn^2+^.

**Figure 6.**
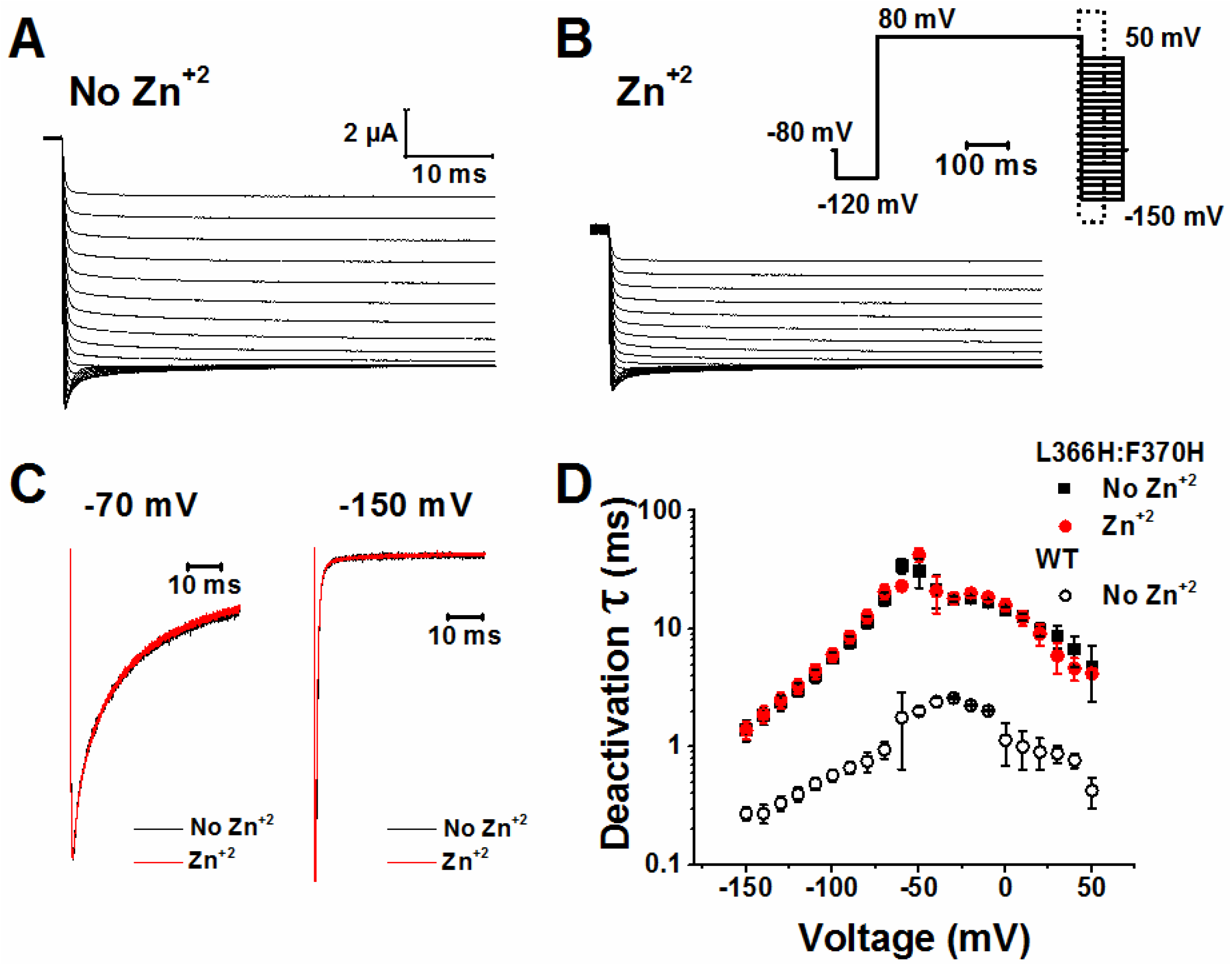
Deactivation time constants of L366H:F370H are not affected by Zn^+2^. **(A-B)** Representative deactivation current traces from an oocyte expressing the mutant L366H:F370H in absence (**A**) and in presence of 20μM Zn^+2^ **(B).** Insert in B is the voltage pulse protocol used to elicit the currents. Dashed rectangle is the time window taken to show the representative currents traces in A and B. **(C)** Two normalized representative ionic currents at different voltages −70 mV (left panel) and −150 mV (right panel) in presence (red line) and absence (black line) of Zn^+2^. **(D)** Deactivation time constant curves for L366H:F370H channel in absence of Zn^+2^ (No Zn^+2^, external solution – filled black squares), and in presence of 20μM Zn^+2^ (Zn^+2^, open red squares). WT time constant in No Zn^+2^ condition is shown as open black circles. Time constants were calculated using eq. 3 and eq. 4. Data are shown as Mean ± SEM (*N* = 7).

These results show that the backward rate, from open (*active*) to closed (*resting* or *intermediate)* state, is not affected by Zn^+2^, indicating a stabilization of a closed state by Zn^+2^ rather than a destabilization of the open state. It is worth mentioning that currents from L366H:F370H mutant channels, in absence of Zn^2+^ and presence of EDTA (to ensure no coordination by heavy metals contamination of the experimental solutions) have slower activation time constant, slower deactivation and its voltage dependence (in steady-state) is rightward shifted compared to WT channels (Fig. 3 A-B and Fig. 6D).

## DISCUSSION

Our functional data provides evidence that Zn^+2^ coordinates two engineered histidine residues, L366H and F370H, in the S4 segment in *Shaker* channels; this coordination is expected to stabilize that particular region of S4 segment in α-helical conformation. The crystallographic-based model of the structure of Kv1.2, a close homologous of *Shaker* potassium channel, shows that the S4 segments are in α-helical conformation in the N-terminal half and 3_10_-helical conformation in the C-terminal half (7). The transition from α-helical to 3_10_-helical occurs between residues V369 and F370 (7). Because the crystal structure of Kv1.2 (homologous of *Shaker* structure) was obtained at zero mV, the conformation is expected to be in the *active*/*relaxed* state. Based on several functional data, it has been proposed that S4 undergoes conformational changes during the activation process (6, 8, 13). Recently, in a quest to show possible transitions between α- and 3_10_-helical conformations in the S4 using Ci-VSP during activation, Kubota et al (2014) used FRET and LRET to measure distances between the two ends of the S4 segments while it transits during the gating process. Those experiments showed no sign of change in the fluorescence signal that could support any conformational change. In addition, it has been suggested that in *Shaker* the hydrophobic plug (a 10 Å thick hydrophobic region contributed by side chains of segments S1 to S3) separates intra from extracellular sides of the voltage sensor domain (7, 9) and stabilizes the S4 region in 3_10_-helical conformation as S4 moves through it during activation (8). Moreover, Infield et al (2018) have used nonsense suppression to investigate the main-chain hydrogen backbone bonding within S4 and they have found that when they perturbed the hydrogen bond at position 369 the *active* conformation of segment S4 was destabilized. Nevertheless, this study did not provide direct evidence of conformational transitions in the S4 segment during its activation process.

The presence of 20μM of Zn^+2^ did not significantly alter the voltage dependence and the activation time constants of K+ currents from WT channels, nor from single mutants L366H or F370H channels (Fig. 2 and 3). We interpret these data, together with the clear effect of Zn^2+^ on L366H:F370H mutants as a strong evidence that there is indeed a coordination between L366H and F370H in the same channel by Zn^+2^. Our study used ionic currents activation, that reflects the last concerted step of the VSD-to-PD electromechanical coupling mechanism. To gain more insight about the conformational changes of the L366H:F370H channel, the study of gating currents (directly related to VSD movement) in this channel would be ideal. However, due to their low expression in oocytes, we were unsuccessful in recording gating currents from this mutant.

If we assume that one His is protonated, it can serve as an organic cation and forms a cation-π interaction with deprotonated His^0^, whereas if His is deprotonated the conjugative π-plane from imidazole side chain can interact with other deprotonated His, by π-π stacking interaction. There is still a third possibility that is when both His are protonated and their interaction is repulsive. Histidine imidazole group in aqueous solution shows a pKa around 6.5. It is unknown whether the His residues are buried into the membrane or exposed to aqueous crevices, therefore pKa of His imidazole group in that region of the VSD is also unknown. Therefore, during our regular experiments at pH 7.4, it is possible that the side chain is either protonated (His^+^) or deprotonated (His^0^). For the pairs that did not show evidence of either Zn^+2^ or pH modulation, (I360H:L366H, I360H:V367H, V363H:L366H, and V363H:V367H, see Fig. S1 and S2), we cannot infer relative to 3_10_-helix stabilization during VSD motion. Possible explanations for this lack of interaction include i) no conformational changes in these regions of S4 are required for activation, and ii) the geometry (orientation of side chains) is not appropriate for His to be coordinated by Zn^+2^. It should be added that, the pH effects on the G-V curves of the channels studied here are consistent with the surface screening charge effects (26–28).

It has been suggested using Ala scanning mutagenesis that C-terminal of S4 segment of *drk1* voltage-gated K^+^ channel presents periodicity consistent with α-helical configuration (29). Moreover, it has been proposed that segment S4 has an *intermediate* state during activation and that S4 requires a short-lived secondary structural change to reach the *active* state (8, 13). Furthermore, by using His scanning mutagenesis, it has been suggested that the S4 segment of *Shaker* exhibits periodicity compatible with 3_10_-helical configuration and that a transition to 3_10_-helical configuration may be necessary to reach the *active* state (13). Our data supports the idea that by stabilizing the α-helical conformation in the region of the third gating charge (R368) of the S4 segment has prevented a normally occurring secondary conformation (3_10_-helix) change during the activation process based on our functional data using the His:His coordinated by Zn^+2^. The stabilization of S4 in α-helical in the presence of Zn^+2^ was enough to dramatically affect the kinetics of activation of L366H:F370H currents (Fig. 3–4), without changing deactivation time constant (Fig. 6) and promoted 10 mV rightward shift in voltage-dependence (Fig. 2 and Table 1). This also implies that closed states (*resting or intermediate)* were stabilized rather than destabilizing the open (*active*) state. Because the deactivation time constant was not affected by Zn^+2^ we infer that Zn^+2^ might not be able to coordinate with the His residues in the *active* state.

Collectively our data suggests a hypothetical and simplified energy landscape for the transitions of the L366H:F370H channel (Fig. 7). We tested this hypothesis with a simple three-state model (*resting, intermediate* and *active* state – Fig. 7A). To simulate the effect of Zn^+2^ in the energy landscape presented in Fig. 7B, we simply divided the rates β and γ by a factor of 20. This makes the *intermediate* energy well deeper. Because we assume that a region of the S4 segment needs a transition to a 3_10_-helical conformation to proceed to the active state, the physical justification of a deeper *intermediate* well in presence of Zn^+2^ is that the α-helical stabilization by Zn^+2^ will make that transition less likely. All the rates used in this model are presented in Fig. S3 (in the support information). This simple model was able to reproduce qualitatively all the major experimental findings. The G-V curve is shifted to the right and its maximum value measured at 1 s is decreased as the Zn^+2^ *(q)* probability of binding to a subunit is increased (Fig. 7C). The model also reproduces the experimentally found increase of the slow component in the time course of the currents (Fig. 7D), and the experimental result of no effect of Zn^+2^ on the deactivation time constants (Fig. 7E-F). A complete description of the model and a complete set of activation currents generated with the model are presented in Fig. S3 (in the Supporting Material). Our simple model was successful to simulate only qualitatively the experimentally recorded currents in presence and absence of Zn^2+^. It is possible that a more complete model could fit the data quantitatively but such a model would need to include all the paths to activation and inactivation of the conduction pathway such as the non-canonical coupling between the voltage sensors and the pore (30, 31).

**Figure 7:**
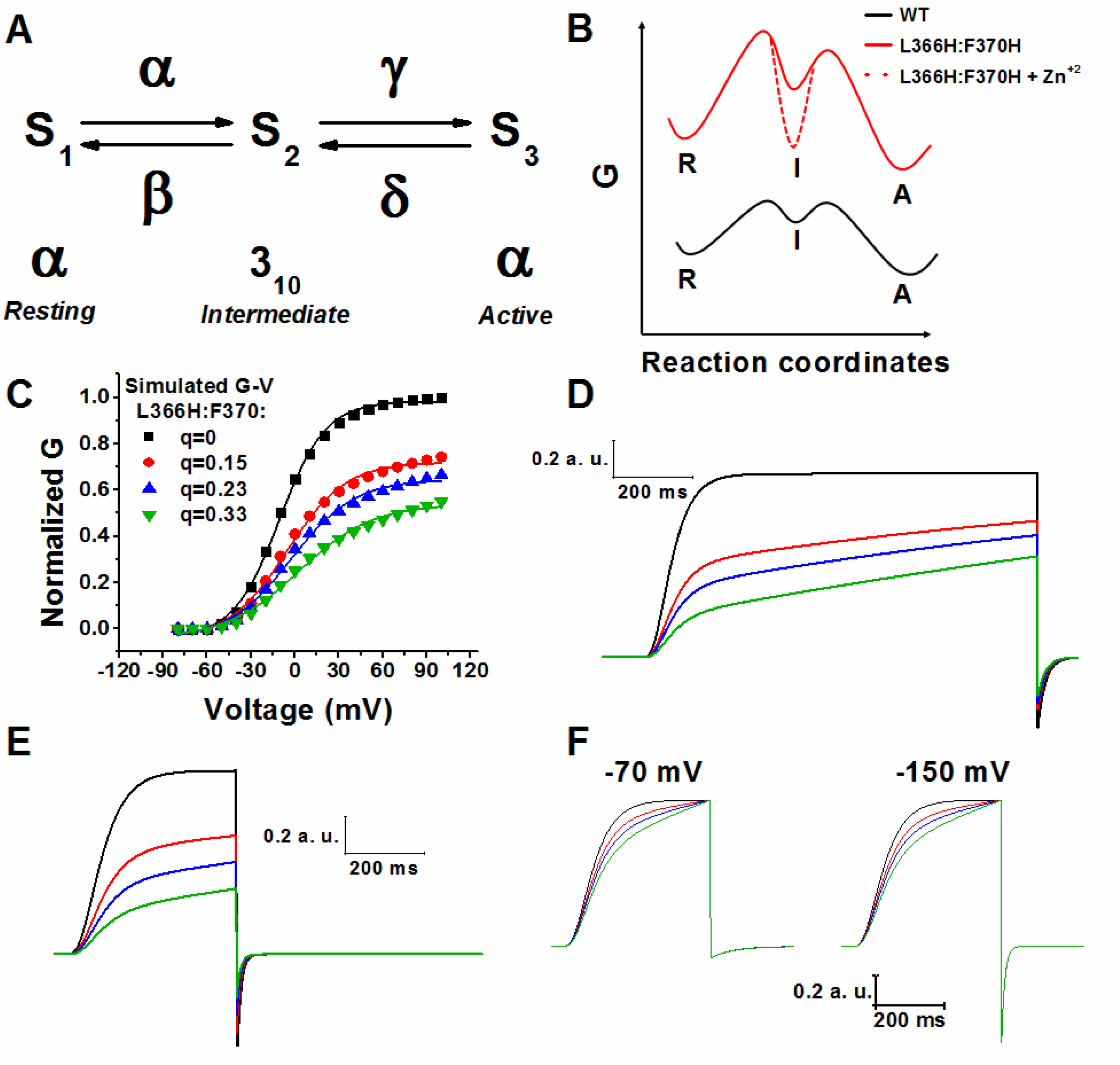
Model of His:His at S4 segment intrasubunit interaction in presence of Zn^+2^ for L366H:F370H. **(A)** Three state model for *Shaker* VSD. *Resting* and *active* states were assumed as α and *intermediate* state as 3_10_-helix. α and β are, respectively, the forward and backward rates for the transition from *resting* to *intermediate* state; γ and δ are, respectively, the forward and backward rates for the transition from *intermediate* to *active* state **(B)** Energy diagram at Vm=0 mV for WT (black), L366:F370H (red) and Zn^+2^ effects on L366H:F370H (dashed red lines). Red dashed line shows that *intermediate* state has the well deepened by Zn^+2^. **C)** Simulated G-V curves calculated using the peak of the currents presented in Fig S3, for L366H:F370H for several Zn^+2^ binding probability q=0 (black squares), q=0.15 (red circles), q=0.23 (blue up triangle) and q=0.33 (green up triangle). G-Vs curves were normalized by the maximum conductance (G) of the channel in absence of Zn^+2^ (*q*=0). Continuous lines over G-V curves are the best fittings of Eq. 2. The fitted parameters are V_1/2_= −10.5 ± 0.7 mV and z= 1.6 ± 0.1 for *q*=0; V_1/2_= −5.2 ± 1.4 mV and z= 1.27 ± 0.1 for *q*=0.15; V_1/2_= −2.7 ± 1.6 mV and z= 1.17 ± 0.1 for *q*=0.23; V_1/2_= 2.7 ± 2 mV and z= 1.0 ± 0.1 for *q*=0.33. **(D)** Simulated activation ionic currents at 100mV. **(E)** Simulated deactivation ionic currents at −150 mV after a 400ms depolarizing pulse of 100mV. **(F)** Two normalized simulated deactivation ionic currents at different voltages −70 mV (left panel) and −150 mV (right panel). The color traces and *q* values used in **D**, **E**, and **F** are the same as shown in **C.**

## CONCLUSION

In summary, we have found that, by stabilizing the S4 segment in α-helical conformation in the region between 366 and 370 of the S4 segment, the activation kinetics is slowed down and the G-V curve is slightly shifted to more depolarized voltages (~ 10mV). This result indicates that α-helical stabilization of this region of the S4 segment prevented a secondary structural short-lived transition. A plausible interpretation is that this *intermediate* state during activation in *Shaker* includes a short-lived 3_10_-helical structure. Although our results cannot be extrapolated to others family of S4-based voltage sensor channels, we believe the strategy of His:His interactions using Zn^+2^ to coordinate them could help to probe transitions in other S4-based voltage sensors which have been structurally and functionally well characterized such as Na^+^ channels and other K^+^ channels.

## AUTHOR CONTRIBUTIONS

Francisco Bezanilla, Carlos A. Z. Bassetto Jr. and Joao Luis Carvalho-de-Souza contributed to the conception and design of the project. CAZBJr performed research and analyzed the data. CAZBJr and JLC-de-S wrote the manuscript with inputs from FB.

## ACKNOWLEDGMENTS

We are thankful to Dr. Bernardo Pinto for helpful discussions and Li Tang for helping with molecular biology. This work was supported by **NIH-R01GM030376 grant.** The authors declare no competing financial interests.

## REFERENCES

1. Hodgkin, A.L., and A.F. Huxley. 1952. A quantitative description of membrane current and its application to conduction and excitation in nerve. J. Physiol. 117: 500–44.

2. Noda, M., S. Shimizu, T. Tanabe, T. Takai, T. Kayano, T. Ikeda, H. Takahashi, H. Nakayama, Y. Kanaoka, N. Minamino, K. Kangawa, H. Matsuo, M.A. Raftery, T. Hirose, S. Inayama, H. Hayashida, T. Miyata, and S. Numa. 1984. Primary structure of Electrophorus electricus sodium channel deduced from cDNA sequence. Nature. 312: 121–127.

3. Long, S.B., E.B. Campbell, and R. Mackinnon. 2005. Crystal structure of a mammalian voltage-dependent Shaker family K+ channel. Science. 309: 897–903.

4. Bezanilla, F. 2000. The Voltage Sensor in Voltage-Dependent Ion Channels. Physiol. Rev. 80: 555–592.

5. Pless, S.A., J.D. Galpin, A.P. Niciforovic, and C.A. Ahern. 2011. Contributions of counter-charge in a potassium channel voltage-sensor domain. Nat. Chem. Biol. 7: 617–623.

6. Vieira-Pires, R.S., and J.H. Morais-Cabral. 2010. 3 10 Helices in Channels and Other Membrane Proteins. J. Gen. Physiol. 136: 585–592.

7. Chen, X., Q. Wang, F. Ni, and J. Ma. 2010. Structure of the full-length Shaker potassium channel Kv1.2 by normal-mode-based X-ray crystallographic refinement. Proc. Natl. Acad. Sci. U. S. A. 107: 11352–7.

8. Henrion, U., J. Renhorn, S.I. Borjesson, E.M. Nelson, C.S. Schwaiger, P. Bjelkmar, B. Wallner, E. Lindahl, and F. Elinder. 2012. Tracking a complete voltage-sensor cycle with metal-ion bridges. Proc. Natl. Acad. Sci. 109: 8552–8557.

9. Lacroix, J.J., H.C. Hyde, F. V. Campos, and F. Bezanilla. 2014. Moving gating charges through the gating pore in a Kv channel voltage sensor. Proc. Natl. Acad. Sci. 111: E1950–E1959.

10. Payandeh, J., T.M. Gamal El-Din, T. Scheuer, N. Zheng, and W.A. Catterall. 2012. Crystal structure of a voltage-gated sodium channel in two potentially inactivated states. Nature. 486: 135–139.

11. Li, Q., S. Wanderling, M. Paduch, D. Medovoy, A. Singharoy, R. Mcgreevy, C.A. Villalba-Galea, R.E. Hulse, B. Roux, K. Schulten, A. Kossiakoff, and E. Perozo. 2014. Structural mechanism of voltage-dependent gating in an isolated voltage-sensing domain. Nat. Struct. Mol. Biol. 21: 244–252.

12. Pan, X., Z. Li, Q. Zhou, H. Shen, K. Wu, X. Huang, J. Chen, J. Zhang, X. Zhu, J. Lei, W. Xiong, H. Gong, B. Xiao, and N. Yan. 2018. Structure of the human voltage-gated sodium channel Nav1.4 in complex with β1. Science. 362: eaau2486.

13. Villalba-Galea, C.A., W. Sandtner, D.M. Starace, and F. Bezanilla. 2008. S4-based voltage sensors have three major conformations. Proc. Natl. Acad. Sci. 105: 17600–17607.

14. Kubota, T., J.J. Lacroix, F. Bezanilla, and A.M. Correa. 2014. Probing α-310 transitions in a voltage-sensing S4 helix. Biophys. J. 107: 1117–1128.

15. Holmgren, M., K.S. Shin, and G. Yellen. 1998. The activation gate of a voltage-gated K+ channel can be trapped in the open state by an intersubunit metal bridge. Neuron. 21: 617–621.

16. Lainé, M., M.C.A. Lin, J.P.A. Bannister, W.R. Silverman, A.F. Mock, B. Roux, and D.M. Papazian. 2003. Atomic proximity between S4 segment and pore domain in shaker potassium channels. Neuron. 39: 467–481.

17. Campos, F. V., B. Chanda, B. Roux, and F. Bezanilla. 2007. Two atomic constraints unambiguously position the S4 segment relative to S1 and S2 segments in the closed state of Shaker K channel. Proc. Natl. Acad. Sci. 104: 7904–7909.

18. Lewis, A., V. Jogini, L. Blachowicz, M. Lainé, and B. Roux. 2008. Atomic constraints between the voltage sensor and the pore domain in a voltage-gated K+ channel of known structure. J. Gen. Physiol. 131: 549–61.

19. Broomand, A., and F. Elinder. 2008. Large-Scale Movement within the Voltage-Sensor Paddle of a Potassium Channel-Support for a Helical-Screw Motion. Neuron. 59: 770–777.

20. Ma, Z., K.Y. Wong, and F.T. Horrigan. 2008. An Extracellular Cu ^2+^ Binding Site in the Voltage Sensor of BK and Shaker Potassium Channels. J. Gen. Physiol. 131: 483–502.

21. Lin, M.A., J.-Y. Hsieh, A.F. Mock, and D.M. Papazian. 2011. R1 in the Shaker S4 occupies the gating charge transfer center in the resting state. J. Gen. Physiol. 138: 155–163.

22. Hoshi, T., W. Zagotta, and R. Aldrich. 1990. Biophysical and molecular mechanisms of Shaker potassium channel inactivation. Science. 250: 533–538.

23. Stefani, E., and F. Bezanilla. 1998. Cut-open oocyte voltage-clamp technique. Methods Enzymol. 293: 300–318.

24. Sundberg, R.J., and R.B. Martin. 1974. Interactions of histidine and other imidazole derivatives with transition metal ions in chemical and biological systems. Chem. Rev. 74: 471–517.

25. Alberts, I.L., K. Nadassy, and S.J. Wodak. 1998. Analysis of zinc binding sites in protein crystal structures. Protein Sci. 7: 1700–1716.

26. Deutsch, C., and S.C. Lee. 1989. Modulation of K+ currents in human lymphocytes by pH. J. Physiol. 413: 399–413.

27. Gilbert, D.L., and G. Ehrenstein. 1969. Effect of Divalent Cations on Potassium Conductance of Squid Axons: Determination of Surface Charge. Biophys. J. 9: 447–463.

28. Gilbert, D.L., and G. Ehrenstein. 1970. Use of a fixed charge model to determine the pK of the negative sites on the external membrane surface. J. Gen. Physiol. 55: 822–825.

29. Li-Smerin, Y., D.H. Hackos, and K.J. Swartz. 2000. alpha-helical structural elements within the voltage-sensing domains of a K(+) channel. J. Gen. Physiol. 115: 33–50.

30. Carvalho-de-Souza, J., and F. Bezanilla. 2017. Non-Canonical Interactions between Voltage Sensors and Pore Domain in Shaker K + -Channel. Biophys. Soc. Meet. Abstr. Biophys.J. 112 (3): Abstract 799-Plat.

31. Fernández-Mariño, A.I., T.J. Harpole, K. Oelstrom, L. Delemotte, and B. Chanda. 2018. Gating interaction maps reveal a noncanonical electromechanical coupling mode in the Shaker K+ channel. Nat. Struct. Mol. Biol. 25: 320–326.

